# Structural and functional conservation of the programmed −1 ribosomal frameshift signal of SARS-CoV-2

**DOI:** 10.1101/2020.03.13.991083

**Authors:** Jamie A. Kelly, Alexandra N. Olson, Krishna Neupane, Sneha Munshi, Josue San Emeterio, Lois Pollack, Michael T. Woodside, Jonathan D. Dinman

**Author notes:** Corresponding authors: Jonathan D. Dinman (mutagenesis and cell-based studies) Michael T. Woodside (*in vitro* MTDB studies) Lois Pollack (SAXS studies).

## Abstract

17 years after the SARS-CoV epidemic, the world is facing the COVID-19 pandemic. COVID-19 is caused by a coronavirus named SARS-CoV-2. Given the most optimistic projections estimating that it will take over a year to develop a vaccine, the best short-term strategy may lie in identifying virus-specific targets for small molecule interventions. All coronaviruses utilize a molecular mechanism called −1 PRF to control the relative expression of their proteins. Prior analyses of SARS-CoV revealed that it employs a structurally unique three-stemmed mRNA pseudoknot to stimulate high rates of −1 PRF, and that it also harbors a −1 PRF attenuation element. Altering −1 PRF activity negatively impacts virus replication, suggesting that this molecular mechanism may be therapeutically targeted. Here we present a comparative analysis of the original SARS-CoV and SARS-CoV-2 frameshift signals. Structural and functional analyses revealed that both elements promote similar rates of −1 PRF and that silent coding mutations in the slippery sites and in all three stems of the pseudoknot strongly ablated −1 PRF activity. The upstream attenuator hairpin activity has also been functionally retained. Small-angle x-ray scattering indicated that the pseudoknots in SARS-CoV and SARS-CoV-2 had the same conformation. Finally, a small molecule previously shown to bind the SARS-CoV pseudoknot and inhibit −1 PRF was similarly effective against −1 PRF in SARS-CoV-2, suggesting that such frameshift inhibitors may provide promising lead compounds to counter the current pandemic.

## Introduction

SARS-CoV2, the etiological agent of COVID-19, is a member of the coronavirus family (1). Coronaviruses have (+) ssRNA genomes that harbor two long open reading frames (ORF) which occupy the 5′ ∼ two-thirds of the genomic RNA (ORF1 and ORF2), followed by several ORFs that are expressed late in the viral replication cycle from subgenomic RNAs (sgRNAs) (**Fig. 1A)** (2). In general, the immediate early proteins encoded by ORF1a are involved in ablating the host cellular innate immune response, whereas the early proteins encoded in ORF1b are involved in genome replication and RNA synthesis. These functions include generating the minus-strand replicative intermediate, new plus-strand genomic RNAs, and subgenomic RNAs which mostly encode structural, late proteins. ORF1b is out of frame with respect to ORF1a, and all coronaviruses utilize a molecular mechanism called programmed −1 ribosomal frameshifting (−1 PRF) as a means to synthesize the ORF2 encoded proteins (3, 4). −1 PRF is a mechanism in which *cis*-acting elements in the mRNA direct elongating ribosomes to shift reading frame by 1 base in the 5′ direction. The use of a −1 PRF mechanism for expression of a viral gene was first identified in the Rous sarcoma virus (5). A −1 PRF mechanism was shown to be required to translate ORF1ab in a coronavirus, Avian Infectious Bronchitis Virus (IBV), two years later (6). In coronaviruses, −1 PRF functions as a developmental switch, and mutations and small molecules that alter this process have deleterious effects on virus replication (7, 8).

**Figure 1.**
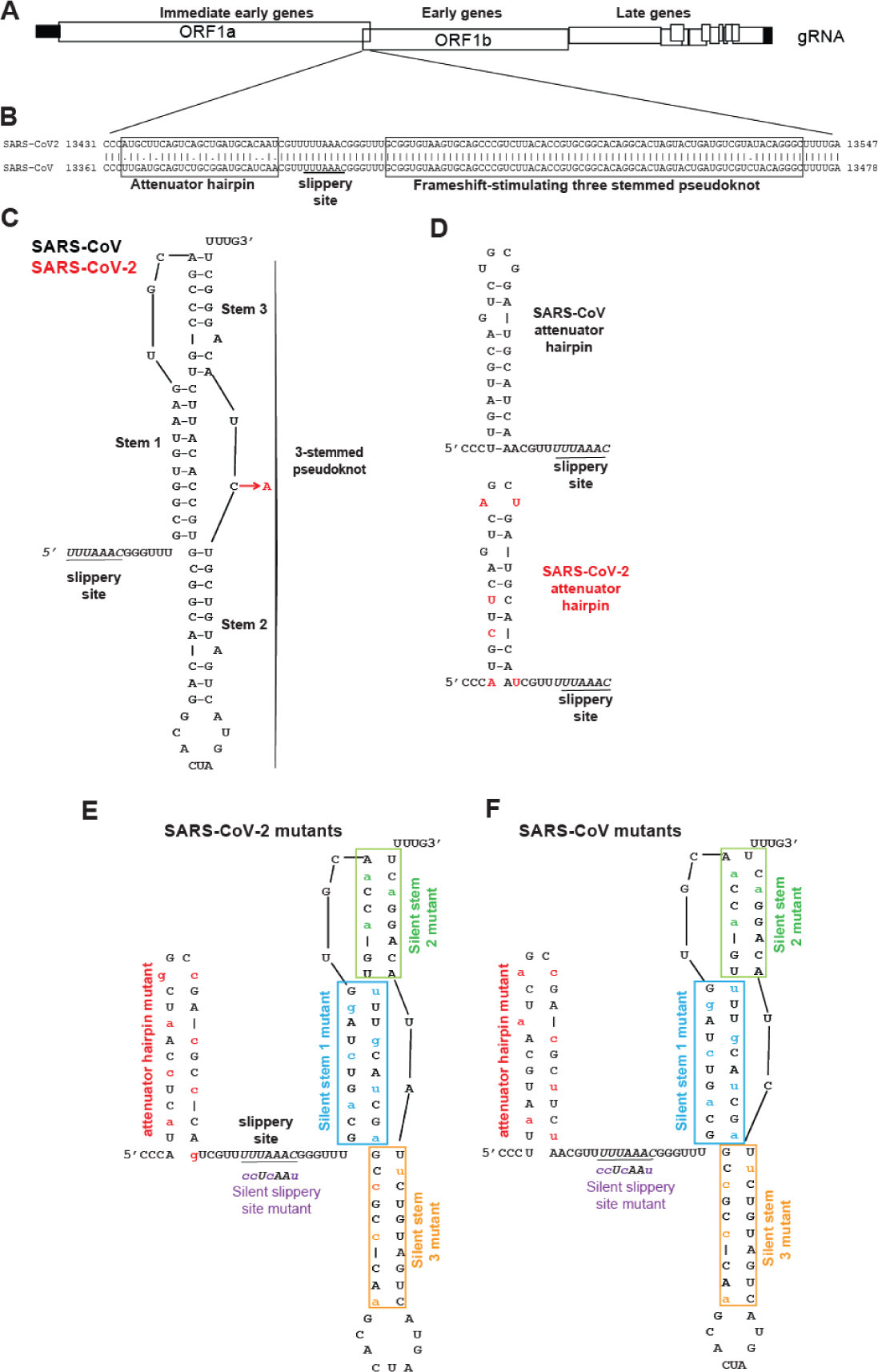
Structural comparison of the SARS-CoV and SARS-CoV-2 −1 PRF signals. **A.** Cartoon depicting SARS-CoV and SARS-CoV2 genome organization including a −1 PRF between ORF1a and ORF1b. **B.** Pairwise analysis of the two −1 PRF signals. The attenuator elements and three-stemmed pseudoknot sequences are boxed as indicated. The U UUA AAC slippery site is underlined. **C.** Structure of the SARS-CoV −1 PRF signal (11) is composed of the 5’ slippery site, a 6-nt spacer, and the three-stemmed pseudoknot stimulatory element. The single base difference in SARS-CoV-2 (red) maps to the short loop linking Stems 2 and 3. **D.** Comparison of the SARS-CoV and SARS-CoV-2 −1 PRF attenuator elements. SARS-CoV-2 specific bases are indicated in red. **E and F.** Silent coding mutations designed to disrupt the attenuators, slippery sites, and Stems 1, 2 and 3 in the SARS-CoV-2 and SARS-CoV −1 PRF signals respectively.

The −1 PRF signal can be broken down into three discrete parts: the “slippery site”, a linker region, and a downstream stimulatory region of mRNA secondary structure, typically an mRNA pseudoknot [reviewed in (3)]. The primary sequence of the slippery site and its placement in relation to the incoming translational reading frame is critical: it must be N NNW WWZ (codons are shown in the incoming or 0-frame), where NNN is a stretch of three identical nucleotides, WWW is either AAA or UUU, and Z ≠ G. The linker region is less well-defined, but typically is short (1 – 12 nt long) and is thought to be important for determining the extent of −1 PRF in a virus-specific manner. The function of the downstream secondary structure is to induce elongating ribosomes to pause, a critical step for efficient −1 PRF to occur [reviewed in (9)]. The generally accepted mechanism of −1 PRF is that the mRNA secondary structure directs elongating ribosomes to pause with its A- and P-site bound aminoacyl- (aa-) and peptidyl-tRNAs positioned over the slippery site. The sequence of the slippery site allows for re-pairing of the tRNAs to the −1 frame codons after they “simultaneously slip” by one base in the 5′ direction along the mRNA. The subsequent resolution of the downstream mRNA secondary structure allows the ribosome to continue elongation of the nascent polypeptide in the new translational reading frame. The downstream stimulatory elements are most commonly H-type mRNA pseudoknots, so called because they are composed of two co-axially stacked stem-loops where the second stem is formed by base pairing between sequence in the loop of the first-stem loop, and additional downstream sequence (10). The SARS-CoV pseudoknot is more complex because it contains a third, internal stem-loop element (11–13). Mutations affecting this structure decreased rates of −1 PRF and had deleterious effects on virus propagation, thus suggesting that it may present a target for small-molecule therapeutics (7, 8). In addition, the presence of a hairpin located immediately 5′ of the slippery site has been reported to regulate −1 PRF by attenuating its activity (14). Here, we report on the −1 PRF signal from SARS-CoV-2. The core −1 PRF signal is nearly identical to that of SARS-CoV, containing only a single nucleotide difference, a C to A. This change maps to a loop region in the molecule that is not predicted to affect the structure of the three-stemmed pseudoknot. The primary sequence of the attenuator hairpin is less-well conserved. However, genetic analyses reveal that both elements appear to have been functionally conserved. Conservation of RNA structure is further supported by the similarity of the small-angle x-ray scattering profiles for the two pseudoknots, and by the similar anti-frameshifting activity of a small-molecule ligand against both frameshift signals.

## Results

### Comparative structural analyses of the two −1 PRF signals

The core of the SARS-CoV −1 PRF signal begins with the U UUA AAC slippery site, followed by a 6-nt spacer region and then the three-stemmed mRNA pseudoknot that stimulates −1 PRF. A second regulatory element, called the attenuator hairpin, is located 5′ of the slippery site. Pairwise analysis of the SARS-CoV and SARS-CoV-2 frameshift signals revealed that the sequence of the attenuator hairpin was less well conserved than the frameshift-stimulating pseudoknot (**Fig. 1B**). The structure of the SARS-CoV −1 PRF signal was previously determined to include a three-stemmed pseudoknot (11). Using this structure as a guide, the single C to A base difference between the core SARS-CoV and SARS-CoV-2 −1 PRF signals (**Fig. 1B**) which maps to a loop that is not predicted to alter the structure of the −1 PRF stimulating element (7) **(Fig. 1C**). In contrast, the attenuator hairpin contains six differences in the nucleotide sequence between the two viruses (**Fig. 1B**), and the SARS-CoV-2 element is predicted to be less stable than its SARS-CoV counterpart (**Fig. 1D**). To determine the importance of each of these elements, a series of silent coding mutants of both the SARS-CoV and SARS-CoV-2 sequences were constructed to disrupt the putative attenuators, slippery sites, and stems 1, 2 and 3 of the pseudoknots (**Fig. 1E, F**).

### Comparative functional analyses of the two −1 PRF signals

Standard dual-luciferase assays were used to monitor −1 PRF activities of the two −1 PRF signals (15, 16) in cultured human cell lines. For both of the elements, −1 PRF activity was ∼20% in HEK (**Fig. 2A**) and ∼30% in HeLa (**Fig. 2B**). Amino acid sequence silent coding mutation of the U UUA AAC slippery sites to C CUC AAC (the incoming 0-frame codons are indicated by spaces) ablated −1 PRF activity in both cases, to less than 1% (**Fig. 2A, B**), demonstrating the functional conservation of this central feature of the −1 PRF signal.

**Figure 2.**
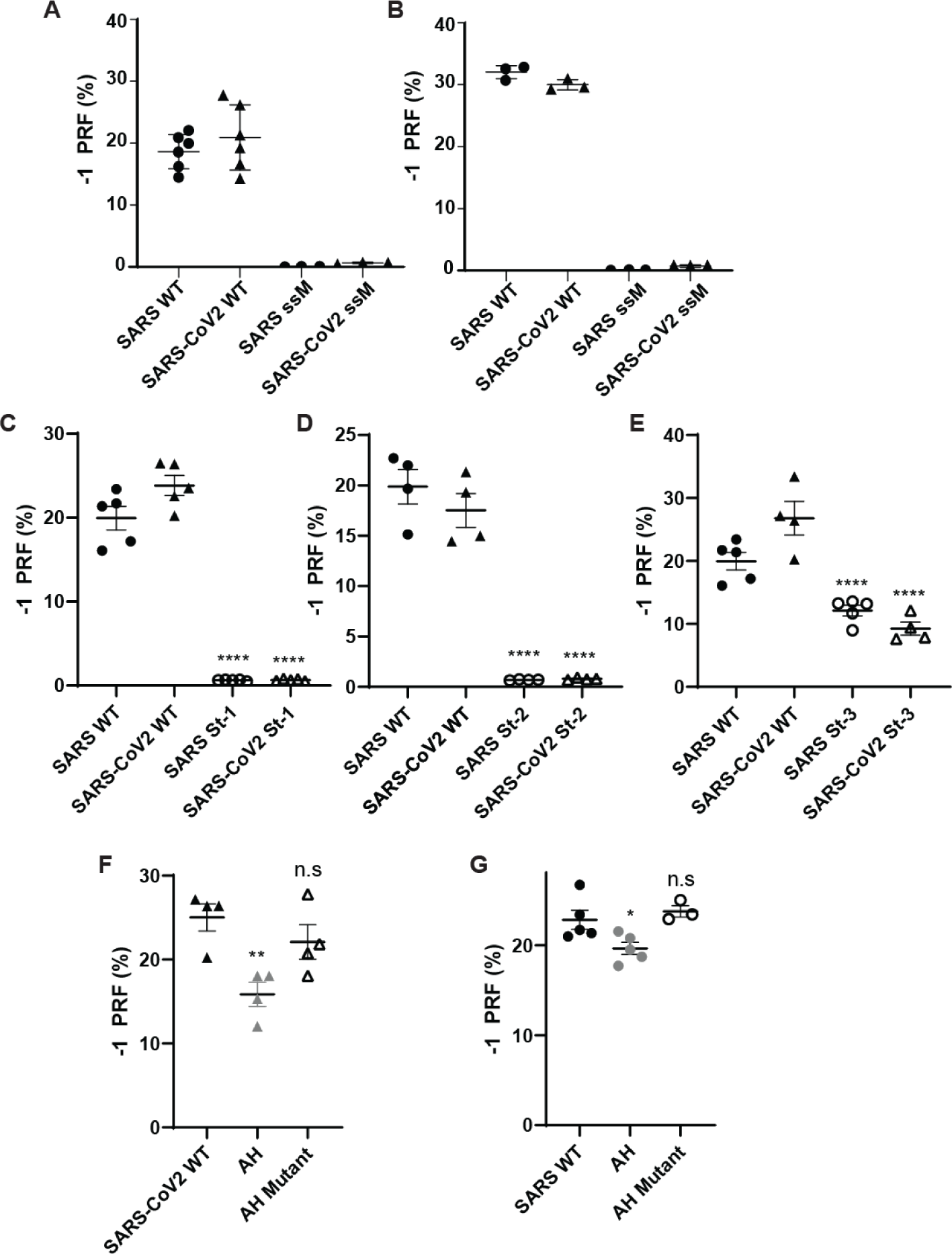
Functional characterization of the SARS-CoV and SARS-CoV-2 −1 PRF signals. **A and B: Analyses of silent slippery site mutants.** The efficiencies of −1 PRF promoted by the wild-type (U UUA AAC) and silent slippery site mutant (C CUC AAC) −1 PRF signals were assayed in HEK (panel A) and HeLA (panel B). ssM denotes silent slippery site mutant. **C – E: Analyses of the importance of the three stems in the -1 PRF stimulating RNA pseudoknot.** Silent stem 1 (St-1, **C**) stem 2 (St-2, **D**) and Stem 3 (St-3, **E**) mutants were assayed in HEK cells. **F and G. Analyses of the attenuator hairpins.** AH denotes constructs that included attenuator hairpin sequences. AH mutant denotes mutants harboring the silent coding attenuator hairpin sequences shown in Figs. 1E, F. Assays were performed using dual-luciferase assays as previously described (15, 16). Each data point represents a single biological replicate comprised of three technical replicates. Error bars denote S.E.M.

To test functional conservation of the three-stemmed pseudoknot, a series of silent 0-frame coding mutations were made to each of the stems in both the SARS-CoV and SARS-CoV-2 frameshift signals and assays were performed in HEK cells. Disruption of Stem 1 strongly suppressed the ability of both elements to promote −1 PRF, decreasing rates to 0.67 ± 0.03% and 0.7 ± 0.1% for SARS-CoV and SARS-CoV-2 respectively, *p* < 0.0001 (**Fig. 2C**). Similarly, disruption of Stem 2 had a strong negative impact on −1 PRF, decreasing rates to 0.68 ± 0.04% for SARS-CoV and 0.8 ± 0.1% for SARS-CoV-2, *p* < 0.0001 (**Fig. 2D**). In contrast, while disruption of Stem 3 did decrease −1 PRF efficiencies, the effects were less severe, although the decreases were statistically significant (13.1 ± 0.9% and 8 ± 1% for SARS-CoV and SARS-CoV-2 respectively, *p* < 0.0001 (**Fig. 2E**). These findings support the hypothesis that structure and function of the core −1 PRF signals has been conserved between the two viruses.

### Conservation of the 5′ attenuator function

Prior studies demonstrated the presence of an element located immediately 5′ of the SARS-CoV slippery site that had the ability to decrease −1 PRF, called the Attenuator Hairpin (14). Although less well conserved at the primary sequence level (**Fig. 1B, C**), addition of this sequence into the SARS-CoV-2 reporter also resulted in decreased −1 PRF efficiency: 16 ± 3% compared to 25 ± 3% without the attenuator hairpin, *p* < 0.01, whereas disruption of the hairpin did not result in decreased efficiency (22 ± 4%, *p* = 0.415) (**Fig. 2F**). In the control experiment, the SARS-CoV attenuator also promoted decreased −1 PRF, albeit to a lesser extent (20 ± 2% compared to 23 ± 2% without the attenuator hairpin, *p* = 0.04, and 24 ± 1% with the disrupted hairpin, *p* = 0.716) (**Fig. 2G**). Thus, the attenuation function has also been conserved between the two viruses despite the differences in primary nucleotide sequences.

### Small-molecule frameshift inhibitor of SARS-CoV −1 PRF is also active against SARS-CoV-2

Based on the strong conservation of the frameshift signal between SARS-CoV and SARS-CoV-2, we tested if a frameshift inhibitor active against the first also retained activity against the second. We focused on a small-molecule ligand previously shown to bind to the SARS-CoV pseudoknot and suppress −1 PRF, 2- {[4-(2-methyl-thiazol-4ylmethyl)- [1,4]diazepane-1-carbonyl]-amino}-benzoic acid ethyl ester, hereafter denoted as MTDB (17, 18). Comparing the −1 PRF activity from dual-luciferase measurements in rabbit reticulocyte lysates in the presence and absence of MTDB, we found that 5 μM MTDB reduced −1 PRF activity by almost 60%, from 36 ± 3% to 15 ± 1% (**Fig. 3**). This reduction was comparable to, but slightly smaller than, that seen previously for the SARS-CoV pseudoknot, where 0.8 μM MTDB reduced −1 PRF by roughly 60% (17).

**Figure 3.**
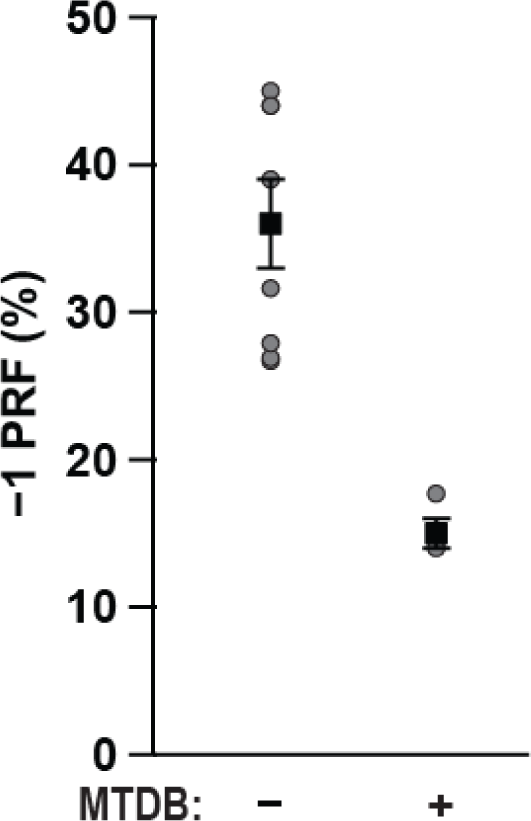
Small-molecule ligand MTDB inhibits −1 PRF stimulation by SARS-CoV-2 pseudoknot. −1 PRF efficiency was reduced almost 60% in the presence of 5 μM MTDB (right), compared to −1 PRF efficiency in the absence of MTDB (left).

### Solution scattering profiles of the SARS-CoV and SARS-CoV-2 pseudoknots are indistinguishable

Finally, we used small and wide-angle x-ray scattering (SAXS) to compare the solution scattering profiles of the two pseudoknots, which reflect their structure. The scattering profiles (intensity as a function of the scattering vector *q*) were indistinguishable for lab purified samples of SARS-CoV (**Fig. 4A, blue**) and SARS-CoV-2 (**Fig. 4A, red**) pseudoknots. The difference between their scattering profiles is consistent with 0 at all *q* (**Fig. 4B**). The high-*q* portion of the profile is sensitive to the finer molecular details of the structure (19) hence the similarity of the profiles for the two pseudoknots indicates their structures are likely the same. Because SARS-CoV pseudoknots can dimerize (20), we also performed inline size exclusion chromatography (SEC) SAXS measurements, where the RNA was purified by SEC immediately before x-ray exposure to ensure only monomers were measured. From inline-SEC SAXS profiles (**Fig. 4A, inset**), we determined the monomer size, parameterized as the radius of gyration, *R*_g_. We measured the same values for SARS-CoV and SARS-CoV-2 pseudoknots: *R*_g_ = 28.1 ± 0.3 Å and *R*_g_ = 28.1 ± 0.2Å, respectively. The difference profile for this set is also consistent with 0 for all *q* (**Fig. 4C**).

**Figure 4.**
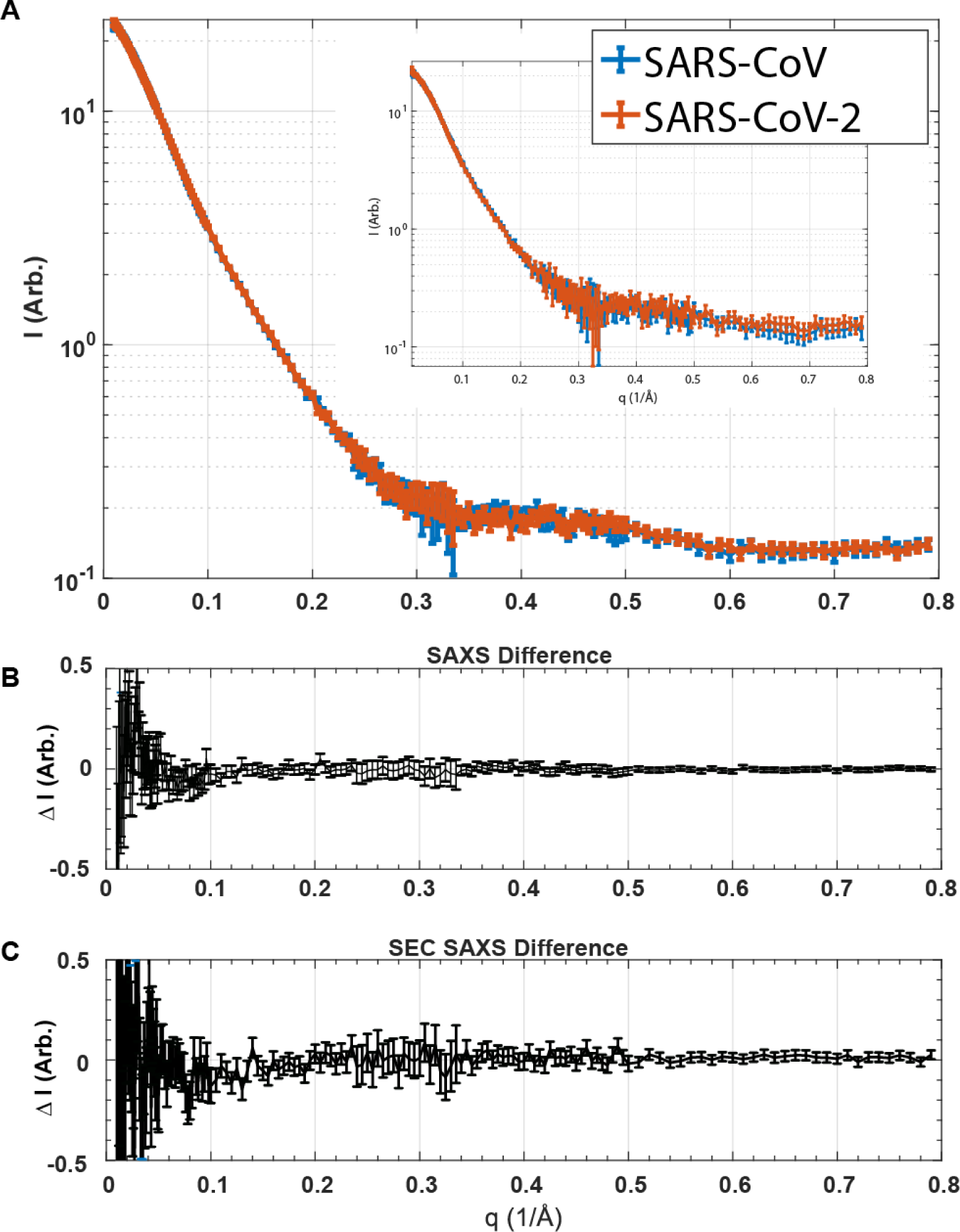
SAXS analyses. (**A**) Scattering profiles from lab purified SAXS samples containing pseudoknots from SARS-CoV (blue) and SARS-CoV-2 (red). Inset: scattering profiles from inline-SEC SAXS measurements, containing purely monomeric pseudoknots. (**B, C**) Difference between the scattering profiles for SARS-CoV and SARS-CoV-2 pseudoknots obtained from lab-purified (**B**) and inline-SEC (**C**) SAXS samples.

## Discussion

These results verify that SARS-CoV-2 does indeed have a functional −1 PRF site. They also show that the properties of the frameshift signal in SARS-CoV-2 are very similar to those of the frameshift signal in SARS-CoV. Not only was the level of −1 PRF close to identical for both viruses, but disrupting stems 1 and 2 in the stimulatory pseudoknot abolished frameshifting in both cases, whereas disrupting stem 3 reduced −1 PRF but did not abolish it in each case. Furthermore, each frameshift signal featured an attenuator hairpin that promoted modestly decreased −1 PRF levels, and the global structures of two pseudoknots as reflected in the SAXS scattering profiles were virtually identical.

The very close correspondence in the properties of the frameshift-stimulatory pseudoknots from SARS-CoV and SARS-CoV-2 suggests that other properties of the former that have been characterized in previous studies are highly likely to carry over to SARS-CoV-2. For example, deletion of stem 3 will likely lead to little or no change in −1 PRF whereas mutation of the A bulge in stem 2 will likely abolish −1 PRF (7, 8), the pseudoknot will likely dimerize via interactions between loop 2 (20), and suppression of −1 PRF will most likely attenuate viral propagation (11). This likely susceptibility of SARS-CoV-2 to attenuation by suppressing −1 PRF is of particular interest, as it suggests that targeting −1 PRF may provide a promising avenue for therapeutic intervention. Previous work on SARS-CoV found that antisense peptide nucleic acids could inhibit both −1 PRF and virus replication (21). The fact that the compound MTDB, which was found in a computational search for −1 PRF inhibitors in SARS-CoV (17), is similarly active at suppressing −1 PRF in SARS-CoV-2 provides concrete evidence for small-molecule frameshifting inhibitors in SARS-CoV-2 and supports the hypothesis that the frameshift-stimulatory pseudoknot may be an attractive therapeutic target.

## Experimental Procedures

### Identification of the SARS-CoV2 −1 PRF signal and computational methods

The SARS-CoV-2 −1 PRF signal was identified from the complete genome sequence (NCBI Ref Seq NC_045512.2). The EMBOSS Water pairwise alignment tool was used to identify sequences in the SARS-CoV-2 genome most similar to the SARS-CoV −1 PRF sequence. One hit was reported between bases 13461 and 13547 of SARS-CoV-2 that was 98.9% identical to the original SARS sequence. The SARS-CoV-2 sequence contains a single point mutation from C to A at base 13533. EMBOSS Water was used to generate pairwise alignments between sequences derived from SARS-CoV (GenBank entry NC_004718.3 begin nt 13361, end nt 13478) and SARS-CoV-2 (Genbank entry NC_045512.2, begin nt 13431, end nt 13547).

### Preparation of plasmids and RNA transcription templates

Plasmids for cell-based dual-luciferase assays for SARS-CoV-2 were generated by site-directed mutagenesis of the pJD2359 plasmid (SARS-CoV pSGDluc reporter plasmid) (8), introducing a single C to A point mutation at base 1873, corresponding to the point mutation in the SARS-CoV-2 genome (Q5 Site-directed mutagenesis kit, NEB). Site directed mutagenesis primers (**Table S1**) were synthesized and purified by IDT. Products were transformed into DH5α Escherichia coli cells (NEB) and spread onto LB agar plates containing 50μg/mL carbenicillin. Positive clones were verified by DNA sequencing (Genewiz). The frameshift reporter negative controls and reporter constructs containing silent mutations disrupting the −1 PRF slippery site (ssM), stem 1 (St1), stem 3 (St3), and attenuator hairpins were constructed similarly by site-directed mutagenesis. Reporters containing silent mutations to stem 2 were made by digesting pJD2257 with SalI and BamHI and ligating a DNA oligonucleotide insert (IDT) containing the silent mutations to stem 2 of SARS and SARS-CoV2 (IDT) into the plasmid using T4 DNA ligase (NEB).

Plasmids for cell-free dual-luciferase assays were made as described previously (22). Briefly, the reporter construct was made by cloning the sequence for Renilla luciferase and SARS-CoV-2 frameshift signal in the 0 frame upstream of the firefly luciferase sequence in the pISO plasmid (Addgene), with firefly luciferase in the −1 frame. A negative control was made by replacing part of the slippery sequence with a stop codon, and a positive control was made without a frameshift signal and the two luciferases in-frame. RNA transcription templates were amplified from these plasmids by PCR and transcribed in vitro by T7 RNA polymerase.

Plasmids for producing samples for SAXS were prepared by ligating an insert containing the sequences of the SARS-CoV and SARS-CoV-2 pseudoknots into the BamHI and SpeI sites of the pMLuc-1 plasmid as described previously (23). RNA transcription templates were amplified from these plasmids by PCR, including 3 extra nucleotides upstream of the pseudoknot and 4 downstream (all U’s). The forward PCR primer was extended on its 5′ end to include the T7 polymerase promoter. The transcription templates were then transcribed in vitro by T7 RNA polymerase.

### Cell culture and plasmid transfection

Human embryonic kidney (HEK293T/17) (CRL-11268) and HeLa (CCL-2) cells were purchased from the American Type Culture Collection (Manassas, VA). HEK293T cells were maintained in Dulbecco’s modified Eagle medium (DMEM) (Fisher Scientific 10-013-CV) supplemented with 10% fetal bovine serum (FBS) (Fisher Scientific 26140-079), 1% GlutaMAX (35050-061), 1% nonessential amino acids (NEAA) (Fisher Scientific 11140-050), 1% HEPES buffer (Fisher Scientific 15630-030) and 1× Penicillin-streptomycin (Fisher Scientific 15140-122) at 37°C in 5% CO_2_. HeLa cells were maintained in DMEM supplemented with 10% FBS, 1% GlutaMAX and 1× Penicillin-streptomycin at 37°C in 5% CO_2_. HEK293T and HeLa cells were seeded at 4 × 10^4^ cells per well into 24-well plates. Cells were transfected 24 hours after seeding with 500 ng dual luciferase reporter plasmid using Lipofectamine3000 (Invitrogen L3000015) per manufacturer’s protocol.

### Dual luciferase assays of −1 PRF

The frameshifting efficiency of the reporter plasmids in cultured cells were assayed as described previously (15, 16) using a dual luciferase reporter assay system kit (Promega). 24 hours post transfection, cells were washed with 1× phosphate-buffered saline (PBS) then lysed with 1× passive lysis buffer (E194A, Promega). Cell lysates were assayed in triplicate in a 96-well plate and luciferase activity was quantified using a GloMax microplate luminometer (Promega). Percent frameshift was calculated by averaging the three firefly or *Renilla* luciferase technical replicate reads per sample then forming a ratio of firefly to *Renilla* luminescence per sample. Each sample luminescence ratio was compared to a read-through control set to 100%. The ratio of ratios for each sample is the percent frameshift for the sample. A minimum of three biological replicates were assayed for each sample, each of which were assayed in triplicate (technical replicates). Mean technical replicate values of each biological replicate are depicted on graphs with standard error of the mean for biological replicates. Statistical analyses were conducted using Student’s *t* test or one-way analysis of variance as appropriate using Prism 8 software (GraphPad).

To measure −1 PRF efficiency in cell-free assays, 2 µg of mRNA from each construct was heated to 65 °C, mixed with 35 µL of nuclease-treated RRL (Promega) and 0.5 µL of 1 mM amino-acid mixture lacking Leu and Met, and then incubated for 90 min at 30°C. The firefly luminescence from each of the constructs was measured after incubating 20 µL of each reaction with 100 µL of Dual-Glo Luciferase reagent (Promega) for 10 min, and then *Renilla* luminescence was measured 10 min after adding 100 µL of Dual-Glo Stop and Glo reagent. The −1 PRF efficiency was calculated from the ratio of firefly and *Renilla* luminescence, subtracting the background measured from the negative control and normalizing by the positive control. Eight independent measurements were made without MTDB, 4 with MTDB.

### SAXS measurements

RNA samples for SAXS experiments were made by in vitro transcription of DNA templates followed by ethanol precipitation of the RNA. To avoid dimerization, RNA was resuspended in a low-salt solution (50mM MOPS 10mM KCl pH 7.5). The RNA was annealed by heating to 95C° for 5 minutes, then placed on ice. After concentration with a spin concentrator, a fraction of the RNA was set aside for inline SEC-SAXS, performed just prior to x-ray exposure, while the rest was purified by size exclusion chromatography in a column equilibrated with the SAXS Buffer (50mM MOPS 130mM KCl pH 7.5). Selected peak fractions of these lab-purified samples were then concentrated to 17.3 μM for the SARS-CoV RNA and 19.2 μM for the SARS-CoV-2 RNA shown in the figure. All samples were sent to the National Synchrotron Light Source II for data acquisition.

SAXS data were collected at the Life Sciences X-Ray Scattering Beamline (LIX) at Brookhaven National Lab using their standard solution scattering set-up, experimental procedures, and data-processing packages (24). SEC-SAXS was performed on a Superdex 200 increase 5/150 GL column (GE) equilibrated in the SAXS buffer condition.

### Data Availability

Full datasets of −1 PRF assays are available upon request. Please contact Dr. Jonathan D. Dinman, University of Maryland.

## Acknowledgements

We wish to thank Kevin Tu for his help with reagent preparation and Shirish Chodankar for support with remote data acquisition at NSLS II. JDD would like to thank the doctors, nurses and staff of the Covid-19 ward at Suburban Hospital for their wonderful care.

