## Supplemental Tables 1 and 2 for "Structural and functional conservation of the programmed −1 ribosomal frameshift signal of SARS-CoV-2"

| **Oligonucleotide Description** | **Sequence** |
| --- | --- |
| SARS-CoV2 −1 PRF WT Fwd | 5’CTGATGTCGTaTACAGGGCTTTTG3’ |
| SARS-CoV2 −1 PRF WT Rev | 5’TACTAGTGCCTGTGCCGC3’ |
| SARS-CoV −1 PRF Attenuator hairpin Fwd | 5’gtctgcggatgcatcaacGTTTTTAAACGGGTTTGCGGTGTAAGTGCAG3’ |
| SARS-CoV −1 PRF Attenuator hairpin Rev | 5’tgcatcaagggttcgcggGTCGACGAGGGCCCGGGG3’ |
| SARS-CoV2 −1 PRF Attenuator hairpin Fwd | 5’gtcagctgatgcacaatcGTTTTTAAACGGGTTTGCGGTGTAAGTGCAG3’ |
| SARS-CoV2 −1 PRF Attenuator hairpin Rev | 5’ tgaagcatgggttcgcggGTCGACGAGGGCCCGGGG 3’ |
| SARS-CoV −1 PRF Attenuator mutant Fwd | 5’ccgacgcttctaCGTTTTTAAACGGGTTTGCGGTGTAAG3’ |
| SARS-CoV −1 PRF Attenuator mutant Rev | 5’ctgattgcattaaGGGTTCGCGGGTCGACGA3’ |
| SARS-CoV2 −1 PRF Attenuator mutant Fwd | 5’ccgacgcccagtCGTTTTTAAACGGGTTTGCGGTGTAAG3’ |
| SARS-Cov2 −1 PRF Attenuator mutant Rev | 5’ccgattggagtatGGGTTCGCGGGTCGACGA3’ |
| SARS-CoV −1 PRF Stem 1 mutant Fwd | 5’cccgttttgcatcgaGCGGCACAGGCACTAGTA3’ |
| SARS-CoV −1 PRF Stem 1 mutant Rev | 5’ctgcacctagactgcAAACCCGTTTAAAAACGTTGATGC3’ |
| SARS-CoV2 −1 PRF Stem 1 mutant Fwd | 5’cccgttttgcatcgaGCGGCACAGGCACTAGTA3’ |
| SARS-CoV2 −1 PRF Stem 1 mutant Rev | 5’ctgcacctagactgcAAACCCGTTTAAAAACGATTGTGC3’ |
| SARS-CoV −1 PRF Stem 3 mutant Fwd | 5’agtactgatgtcttCTACAGGGCTTTTGATGGATC3’ |
| SARS-CoV −1 PRF Stem 3 mutant Rev | 5’agtgcttgggcggcACGGTGTAAGACGGGCTG3’ |
| SARS-CoV2 −1 PRF Stem 3 mutant Fwd | 5’agtactgatgtcttATACAGGGCTTTTGATGGATCC3’ |
| SARS-CoV2 −1 PRF Stem 3 mutant Rev | 5’agtgcttgggcggcACGGTGTAAGACGGGCTG3’ |
| SARS-CoV −1 PRF Stem 2 mutant Top | 5’tcgacgTTTTTAAACGGGTTTGCGGTGTAAGTGCAaCCaGTCTTACACCGTGCGGCACAGGCACTAGTACTGATGTCGTCTACAGGaCTTTTGATg 3’ |
| SARS-CoV −1 PRF Stem 2 mutant Bottom | 5’gatccATCAAAAGtCCTGTAGACGACATCAGTACTAGTGCCTGTGCCGCACGGTGTAAGACtGGtTGCACTTACACCGCAAACCCGTTTAAAAAcg 3’ |
| SARS-CoV2 −1 PRF Stem 2 mutant Top | 5’tcgacgTTTTTAAACGGGTTTGCGGTGTAAGTGCAaCCaGTCTTACACCGTGCGGCACAGGCACTAGTACTGATGTCGTATACAGGaCTTTTGATg 3’ |
| SARS-CoV2 −1 PRF Stem 2 mutant Bottom | 5’gatccATCAAAAGtCCTGTATACGACATCAGTACTAGTGCCTGTGCCGCACGGTGTAAGACtGGtTGCACTTACACCGCAAACCCGTTTAAAAAcg 3’ |
| SARS-CoV2 −1 PRF FWD | 5’gTTTTTAAACGGGTTTGCGGTGTAAGTGCAGCCCGTCTTACACCGTGCGGCACAGGCACTAGTACTGATGTCGTATACAGGGCTTTa 3’ |
| SARS-CoV2 −1 PRF REV | 5’ctagtAAAGCCCTGTATACGACATCAGTACTAGTGCCTGTGCCGCACGGTGTAAGACGGGCTGCACTTACACCGCAAACCCGTTTAAAAActgca 3’ |
| SARS-CoV2 0 PRF FWD | 5’gTTTAGAAACGGGTTTTGCGGTGTAAGTGCAGCCCGTCTTACACCGTGCGGCACAGGCACTAGTACTGATGTCGTATACAGGGCTTTa 3’ |
| SARS-CoV2 0 PRF REV | 5’ctagtAAAGCCCTGTATACGACATCAGTACTAGTGCCTGTGCCGCACGGTGTAAGACGGGCTGCACTTACACCGCAAAACCCGTTTCTAAActgca 3’ |
| SARS-CoV2 STOP FWD | 5’gTTTTGAAACGGGTTTGCGGTGTAAGTGCAGCCCGTCTTACACCGTGCGGCACAGGCACTAGTACTGATGTCGTATACAGGGCTTTTa 3’ |
| SARS-CoV2 STOP REV | 5’ctagtAAAAGCCCTGTATACGACATCAGTACTAGTGCCTGTGCCGCACGGTGTAAGACGGGCTGCACTTACACCGCAAACCCGTTTCAAAActgca 3’ |

**Supplementary Table 1. Synthetic oligonucleotides used in this study.**

| **Plasmid** | **Description** |
| --- | --- |
| pJD2257 | Modified pSGDluc (dual luciferase with inteins) Readthrough control |
| pJD2359 | Modified pSGDluc (dual luciferase with inteins) with SARS-CoV −1 PRF insert |
| pJD2514 | Modified pSGDluc (dual luciferase with inteins) with SARS-CoV-2 −1 PRF insert |
| pJD2515 | Modified pSGDluc (dual luciferase with inteins) with SARS-CoV silent slippery site mutant insert |
| pJD2516 | Modified pSGDluc (dual luciferase with inteins) with SARS-CoV-2 silent slippery site mutant insert |
| pJD2517 | Modified pSGDluc (dual luciferase with inteins) with SARS-CoV attenuator hairpin and −1 PRF insert |
| pJD2518 | Modified pSGDluc (dual luciferase with inteins) with SARS-CoV disrupted attenuator hairpin and −1 PRF insert |
| pJD2519 | Modified pSGDluc (dual luciferase with inteins) with SARS-CoV2 attenuator hairpin and −1 PRF insert |
| pJD2520 | Modified pSGDluc (dual luciferase with inteins) with SARS-CoV-2 disrupted attenuator hairpin and −1 PRF insert |
| pJD2521 | Modified pSGDluc (dual luciferase with inteins) with SARS-CoV −1 PRF insert containing silent mutations in Stem 1 |
| pJD2522 | Modified pSGDluc (dual luciferase with inteins) with SARS-CoV −1 PRF insert containing silent mutations in Stem 2 |
| pJD2523 | Modified pSGDluc (dual luciferase with inteins) with SARS-CoV −1 PRF insert containing silent mutations in Stem 3 |
| pJD2524 | Modified pSGDluc (dual luciferase with inteins) with SARS-CoV-2 −1 PRF insert containing silent mutations in Stem 1 |
| pJD2525 | Modified pSGDluc (dual luciferase with inteins) with SARS-CoV-2 −1 PRF insert containing silent mutations in Stem 2 |
| pJD2526 | Modified pSGDluc (dual luciferase with inteins) with SARS-CoV-2 −1PRF insert containing silent mutations in Stem 3 |
| DTO20 | pISO plasmid modified with insert containing *Renilla* luciferase and pMLuc1 multiple cloning site |
| DTO20-1FSWu | DTO20 with SARS-CoV-2 −1PRF insert |
| DTO20-0FSWu | DTO with SARS-CoV-2 0 PRF insert |
| DTO20-STOPWu | DTO with SARS-CoV-2 −1PRF insert modified with stop codon in the slippery sequence |

**Supplementary Table 2. Plasmids used in this study.**
